# Prohibitin depletion extends lifespan of a TORC2/SGK-1 mutant through autophagy

**DOI:** 10.1101/792465

**Authors:** Blanca Hernando-Rodríguez, Mercedes M. Pérez-Jiménez, María Jesús Rodríguez-Palero, Antoni Pla, Manuel David Martínez-Bueno, Patricia de la Cruz Ruiz, Roxani Gatsi, Marta Artal-Sanz

## Abstract

Mitochondrial prohibitins (PHB) are highly conserved proteins with a peculiar effect on lifespan. While PHB depletion shortens lifespan of wild type animals, it enhances longevity of a plethora of metabolically compromised mutants, including target of rapamycin complex 2 (TORC2) mutants *sgk-1* and *rict-1*. Here, we show that *sgk-1* mutants have impaired mitochondrial homeostasis, lipogenesis, yolk formation and autophagy flux due to alterations in membrane lipid and sterol homeostasis. Remarkably, all these features are suppressed by PHB depletion. Lifespan analysis shows that autophagy and the mitochondrial unfolded protein response (UPR^mt^), but not mitophagy, are required for the enhanced longevity caused by PHB depletion in *sgk-1* mutants. We hypothesize that UPR^mt^ induction upon PHB depletion extends lifespan of *sgk-1* mutants through autophagy. Our results strongly suggest that PHB depletion suppresses the autophagy defects of *sgk-1* mutants by altering membrane lipid composition at ER-mitochondria contact sites, where TORC2 localizes.

## Introduction

Mitochondrial function, nutrient signalling and autophagy regulate aging across phyla. However, their exact mechanisms and interactions in lifespan modulation still remain elusive. The mitochondrial prohibitin (PHB) complex is a strongly evolutionarily conserved ring-like macromolecular structure (Artal-Sanz and Tavernarakis, 2009a; Merkwirth and Langer, 2009). While deletion of PHB does not cause any observable growth phenotype in the unicellular yeast *Saccharomyces cerevisiae* (Osman et al., 2009), in *Caenorhabditis elegans* the PHB complex is required for embryonic development and its postembryonic depletion leads to morphological abnormalities in the somatic gonad and sterility (Artal-Sanz et al., 2003). Prohibitins have been implicated in several age-related diseases (Nijtmans et al., 2002; Thuaud et al., 2013) and are involved in mitochondrial morphogenesis and membrane maintenance by acting as chaperones (Nijtmans et al., 2000) and scaffolds (Merkwirth et al., 2012). PHB depletion has been shown to perturb mitochondrial homeostasis causing an induction of the mitochondrial unfolded protein response, UPR^mt^ (Benedetti et al., 2006; Hernando-Rodriguez et al., 2018; Yoneda et al., 2004).

PHB depletion has opposing effects on lifespan depending on the genetic background. Loss of PHB by RNAi shortens lifespan in wild type worms, whereas it increases lifespan in different metabolically compromised backgrounds. These include mutants of the Insulin/IGF-1 signalling (IIS) pathway and mutants with compromised mitochondrial function or fat metabolism (Artal-Sanz and Tavernarakis, 2009b). In particular, PHB depletion increases lifespan of the long-lived *daf-2(e1370)* mutants, where the induction of the UPR^mt^ is reduced (Gatsi et al., 2014). Analysis of known kinases acting downstream of the insulin receptor DAF-2 revealed that loss of function of SGK-1 in PHB depleted animals results in enhanced longevity and reduced UPR^mt^ activation similar to that observed in PHB depleted *daf-2* mutants (Gatsi et al., 2014). SGK-1 belongs to the AGC kinase family and is the sole *C. elegans* homologue of the mammalian Serum- and Glucocorticoid-inducible Kinase. Its activation depends on PI3K and has been involved in response to oxidative stress and DNA damage, among others (Di Cristofano, 2017). In addition to acting in the IIS pathway, SGK-1 regulates aging and mitochondrial homeostasis through a parallel pathway, as part of TORC2 (Target Of Rapamycin Complex 2), downstream of RICT-1. Interestingly, depletion of PHB in *daf-2;sgk-1* double mutants enhances both the increased lifespan and the reduction of the UPR^mt^ (Gatsi et al., 2014).

Work in yeast has demonstrated that Ypk1, the yeast homologue of SGK-1, is the relevant target of TORC2. Ypk1 is involved in sphingolipid synthesis and ceramide signalling, having an essential role in lipid membrane homeostasis and affecting cell size and growth rate (Lucena et al., 2018; Muir et al., 2014; Roelants et al., 2018). TORC2-Ypk1 controls sphingolipid homeostasis by sensing and regulating ROS (Niles et al., 2014). Moreover, TORC2-Ypk1 regulates autophagy induced by amino acid starvation (Vlahakis et al., 2014) and modulates the autophagy flux by controlling mitochondrial respiration and calcium signalling (Vlahakis et al., 2016; Vlahakis et al., 2017). The underlying molecular mechanism remains however unknown. In worms, SGK-1 regulates development, fat metabolism, stress responses and lifespan in a complex and controversial manner, partially explained by the different alleles under study and the different growing conditions (Chen et al., 2013; Dowen et al., 2016; Gatsi et al., 2014; Hertweck et al., 2004; Jones et al., 2009; Mizunuma et al., 2014; Soukas et al., 2009; Tullet et al., 2008; Zhou et al., 2019).

To better understand how SGK-1 regulates lifespan and mitochondrial function, we explored the interaction between PHB and SGK-1. Our results show that long-lived *sgk-1(ok538)* mutants have altered mitochondrial structure and function, phenotypes that are suppressed by PHB depletion. A transcription factor RNAi screen identified membrane lipid homeostasis as a mechanism implicated in SGK-1 mediated maintenance of mitochondrial function. In agreement with that, electron microscopy analysis showed defects in organelle membrane contact sites in *sgk-1* mutants, leading to defective lipogenesis and lipoprotein production and a blockage of the autophagic flux. Remarkably, these phenotypes are suppressed by PHB depletion. Furthermore, lifespan analyses showed that autophagy and the UPR^mt^ are required for the enhanced longevity of *sgk-1* mutants upon PHB depletion, while mitophagy is not. Consequently, in the absence of the PHB complex, SGK-1 expression levels increased upon inhibition of the UPR^mt^, but are not affected by inhibition of mitophagy. Our data strongly suggest that UPR^mt^ induction upon PHB depletion extends lifespan of *sgk-1* mutants through autophagy. We discuss our observations in light of the conserved role of TORC2 in lipid membrane biology and the proposed role of the PHB complex and TORC2 at mitochondria-associated endoplasmic reticulum (ER) membranes (MAM).

## Results

### Prohibitin depletion suppresses the altered mitochondrial structure and function of *sgk-1* mutants

In *C. elegans*, loss of function of SGK-1 causes a delay in development (Jones et al., 2009) as well as reduced brood and body size (Soukas et al., 2009). These phenotypes are consistent with phenotypes observed in mitochondrial mutants (Dillin et al., 2002). In addition, worms lacking the TORC2 component RICT-1 or the downstream kinase SGK-1, have an induced mitochondrial unfolded protein response (UPR^mt^) (Gatsi et al., 2014). We analyzed whether mitophagy, another mitochondrial quality control mechanism, might be activated in *sgk-1* deletion mutants using a mitophagy reporter based on PINK-1 protein (Kirienko et al., 2015). PINK-1 is a serine threonine protein kinase that, when mitochondria are depolarized, accumulates in the outer mitochondrial membrane and targets mitochondria for selective autophagy (Narendra and Youle, 2011; Palikaras et al., 2015). We observed that depletion of *sgk-1* increased PINK-1 protein levels (Figure 1A). Similarly, PINK-1 protein levels increased upon depletion of *phb-1* or *atfs-1,* the key UPR^mt^ regulator (Figure 1A), confirming that *sgk-*1 mutants suffer from mitochondrial stress.

**Figure 1:**
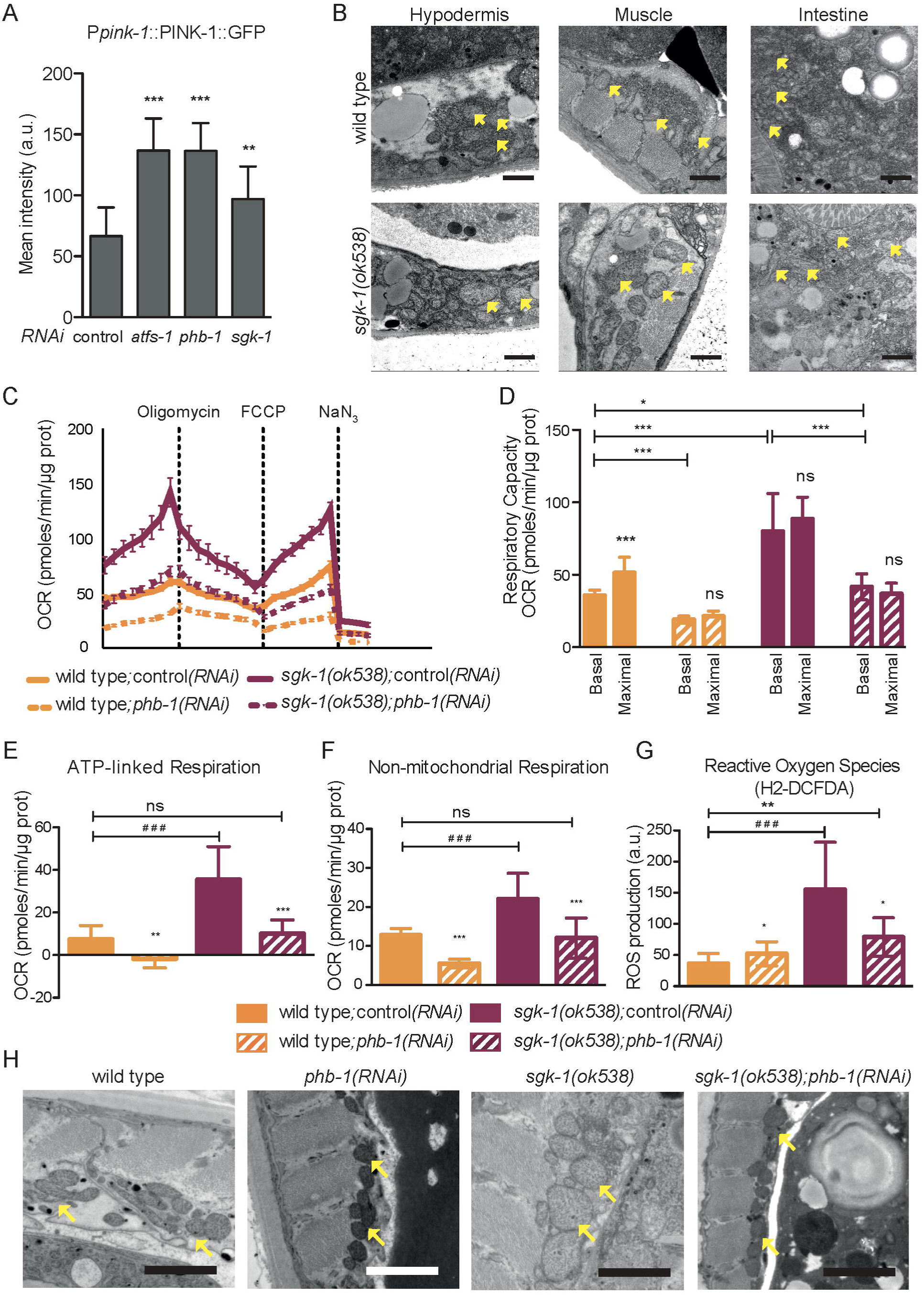
Prohibitin depletion suppresses the altered mitochondrial structure and function of *sgk-1* mutants. A. Quantification of Ppink-1::PINK-1::GFP in worms treated with RNAi against *atfs-1, phb-1* and *sgk-1* at the young adult stage. **B.** Electron microscopy images of hypodermis, muscle and intestine of wild type and *sgk-1* mutants at day 1 of adulthood. Yellow arrows show mitochondria. Bar size: 1 µm. **C-F.** Seahorse measurements in wild type animals, *phb-1(RNAi)* treated worms, *sgk-1* mutants and *sgk-1;phb-1* depleted mutants at the young adult stage. **C.** Mitochondrial oxygen consumption rate. **D.** Basal and maximal respiratory capacity. **E.** ATP-linked respiration and **F.** Non-mitochondrial respiration. **G.** Reactive oxygen species (ROS) levels measured at the young adult stage of wild type animals, *phb-1(RNAi)* treated worms, *sgk-1* mutants and *sgk-1;phb-1* depleted mutants. **H.** Electron microscopy images of muscle cells, at day 5 of adulthood, of wild type and *sgk-1(ok538)* mutants under normal and mitochondrial stress conditions *(phb-1(RNAi)).* Yellow arrows point to mitochondria. Bar size: 1 µm. (Mean ± SD; *** *p* value < 0.001, ** *p* value < 0.01, * *p* value < 0.1, ns not significant compared against its respective *control(RNAi),* ### *p* value < 0.001 compared against wild type *control(RNAi); t-test.* Combination of three independent replicates).

To better define the mitochondrial defect of *sgk-1* deletion mutants we performed transmission electron microscopy (TEM) analysis. Mitochondria in *sgk-1* mutants were swollen, and bigger compared to wild type worms in all tissues analyzed (hypodermis, muscle and intestine) at both, day one and day five of adulthood (Figure 1B and Figure S1A, respectively). In contrast to previous observations (Zhou et al., 2019), no obvious mitochondrial fragmentation or severe cristae defects in the intestine were observed in the TEM sections analyzed.

To better understand the interaction between PHB and SGK-1, we measured mitochondrial performance in *sgk-1* mutants in the presence or absence of the PHB complex, during aging (Figures 1C-F and Figure S1B-C). At the young adult stage, *sgk-1* mutants showed a dramatic increase in basal OCR compared to wild type worms (Figures 1C and 1D), consistent with the previous finding that mammalian mTORC2/rictor knockdown leads to an increase in mitochondrial respiration in mammalian cells (Schieke et al., 2006). The increased respiration rate of *sgk-1* mutants is suppressed by PHB depletion, which also reduces the OCR of wild type young adult worms (Figures 1C and 1D). At day 6 of adulthood, the OCR was reduced compared to young animals (Figure S1C) as described (Van Raamsdonk et al., 2010), but not in *phb-1(RNAi)* worms (Figure S1C). In *sgk-1* mutants, respiration dramatically drops during aging, to the level of wild type animals. However, PHB depletion increases the OCR in wild type and *sgk-1* mutants at day 6 of adulthood (Figure S1B and S1C). This suggests that mitochondrial function declines with age in wild type animals but this decline occurs at a higher rate in *sgk-1* mutants. In contrast, respiration remains relatively constant in PHB depleted worms, where respiration is compromised already at the young adult stage.

Measurements of maximal respiratory capacity at the young adult stage showed that *sgk-1* mutants had a higher maximal respiratory capacity compared to wild type worms, which was fully suppressed by PHB depletion. PHB depletion also reduced the maximal respiratory capacity of wild type worms (Figure 1D). The spare capacity, is the difference between maximal and basal OCR and it is defined as the amount of oxygen that is available for cells under stress or increased energy demands. *sgk-1* mutants showed an insignificant spare respiratory capacity compared to wild type animals. Additionally, depletion of *phb-1* reduced the spare capacity of both *sgk-1* and wild type animals (Figure 1D). We observed increased ATP-linked OCR in *sgk-1* mutants (Figure 1E), which may correspond to a higher energy demand. *phb-1(RNAi)* reduced the ATP-linked OCR (Figure 1E) both in otherwise wild type worms and in *sgk-1* mutants, suggesting that the oxidative phosphorylation system was severely compromised. Again, depletion of the PHB complex reversed *sgk-1* mutant phenotype, bringing ATP-linked OCR to wild-type levels (Figure 1E). The O_2_ consumed after sodium azide (NaN_3_) injection corresponds to the respiration caused by other cellular processes than oxidative phosphorylation, such as non-mitochondrial NADPH oxidases or reactive oxygen species production. Interestingly, *sgk-1* mutants showed an increased non-mitochondrial respiration compared to wild type at the young adult stage, which was suppressed by PHB depletion. Lack of PHB also reduced non-mitochondrial respiration in wild type worms (Figure 1F).

Mitochondrial respiration is the predominant source of cellular ROS. We thus evaluated ROS levels in *sgk-1* mutants in the presence and absence of the PHB complex. We measured ROS formation in young adults using the dye H2-DCFDA (Figure 1G). In agreement with the observed increase in both, mitochondrial and non-mitochondrial respiration, *sgk-1* mutants showed dramatically elevated ROS levels in comparison to wild type worms, as previously published in other organisms (Niles et al., 2014; Wang et al., 2019). Depletion of *phb-1* increased ROS levels in wild type worms (Figure 1G and (Artal-Sanz and Tavernarakis, 2009b)). However, depletion of *phb-1* reduced ROS levels in *sgk-1* mutants (Figure 1G). Since H2-DCFDA measures mostly cytosolic ROS, these results indicate that *sgk-1* mutants have high levels of cytosolic ROS that are reduced upon *phb-1* depletion. Since increased mitochondrial ROS extends lifespan of mitochondrial mutants with high levels of cytoplasmic ROS (Schaar et al., 2015), we wondered whether the enhancement of lifespan upon *phb-1* depletion in *sgk-1* mutants may be due to an increase in mitochondrial ROS levels. Absence of cytosolic superoxide dismutase, *sod-1*, did not affect lifespan of wild type worms, while lack of the mitochondrial superoxide dismutase, *sod-2*, increased wild type lifespan (Figure S1D, Table S1 and (Van Raamsdonk and Hekimi, 2009; Yang and Hekimi, 2010)). Interestingly, lack of *sod-1* reduced lifespan of *sgk-1* mutants, indicating that increased cytosolic ROS is detrimental for *sgk-1* lifespan, while depletion of *sod-2*, increased lifespan of *sgk-1* mutants (Figure S1E, Table S1), similar to mitochondrial mutants (Schaar et al., 2015). In order to test if mitochondrial ROS generation upon PHB depletion extends lifespan of *sgk-1* mutants, we treated animals with the antioxidant N-acetyl-cysteine (NAC). NAC did not affect the lifespan of wild type worms or *sgk-1* mutants. However, it shortened that of *phb-1* depleted backgrounds, both wild type and *sgk-1* mutants (Figure S1F and S1G, Table S1). Therefore, we cannot conclude that ROS is responsible for the lifespan extension conferred by PHB depletion to *sgk-1* mutants.

Finally, we wondered whether depletion of the PHB complex would suppress the increased mitochondrial size observed in *sgk-1* mutants. TEM analysis of muscle mitochondria showed that *phb-1* depletion drastically reduced mitochondrial size in *sgk-1* mutants, and resulted in mitochondrial fragmentation as it happens in otherwise wild type worms (Artal-Sanz et al., 2003) (Figure 1H).

### Altered membrane sterol and lipid homeostasis induces the UPR^mt^ in the TORC2/SGK-1 mutant

In order to understand the interaction of PHB with SGK-1 in lifespan regulation, we looked for transcription factors mediating the UPR^mt^ in *sgk-1* mutants, as they might be implicated in the activation of pro-survival pathways responsible for the long lifespan observed in PHB-depleted TORC2 mutants (Gatsi et al., 2014). Several transcription factors have been involved in the maintenance of mitochondrial homeostasis and lifespan, including DAF-16, SKN-1, HIF-1 and HSF-1 (Fang et al., 2017; Hwang et al., 2014; Kamal et al., 2016; Labbadia et al., 2017; Lee et al., 2010; Mouchiroud et al., 2013; Palikaras et al., 2015; Senchuk et al., 2018; Tullet et al., 2008; Wu et al., 2018b). Moreover, the UPR^mt^ promotes longevity by activating HIF-1, DAF-16 and SKN-1 (Wu et al., 2018b). We investigated the involvement of these transcription factors in the activation of the UPR^mt^ in *sgk-1* mutants and the essential TORC2 component *rict-1.* Lack of DAF-16 slightly induced mitochondrial stress in wild type animals, while only HIF-1 depletion futher induced the UPR^mt^ in *sgk-1* mutants (Figure 2A). However, in the *rict-1* mutant, depletion of all the transcription factors enhanced the UPR^mt^ (Figure S2A). This suggests that TORC2 deficiency modulates the UPR^mt^ through other kinases in addition to *sgk-1*. Nevertheless, depletion of none of the transcription factors tested inhibited the expression of *hsp-6*, suggesting that different mechanisms are implicated in the UPR^mt^ in TORC2/SGK-1 mutants.

**Figure 2:**
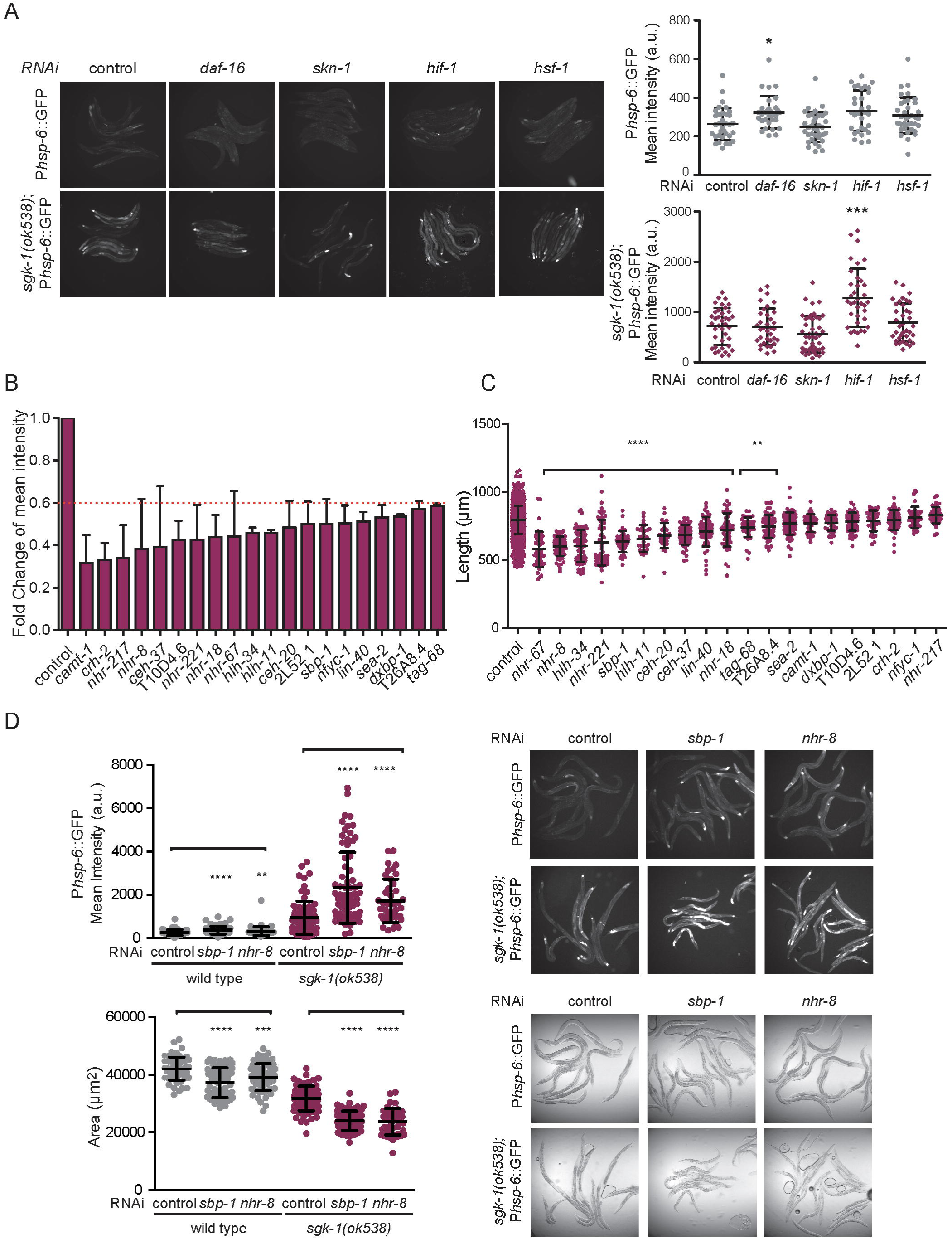
Altered membrane sterol and lipid homeostasis induces the UPR^mt^ in the TORC2/SGK-1. **A.** Quantification of the UPR^mt^ reporter *phsp-6*::GFP in wild type animals (top) and *sgk-1* (bottom) upon depletion of *daf-16, skn-1, hif-1* and *hsf-1.* (Mean ± SD; *** *p* value < 0.001, * *p* value < 0.1 compared against control(RNAi); ANOVA test). Representative images are shown. **B.** Fold change of GFP intensity of the 20 clones whose depletion reduces the mitochondrial stress response in *sgk-1* mutants. **C.** Length of the worms (µm) upon depletion of the 20 candidates in *sgk-1* mutants. **D.** Quantification of the UPR^mt^ reporter (left upper panel) and worm area (left bottom panel) in wild type animals and *sgk-1* mutants upon depletion of *sbp-1* and *nhr-8* (Mean ± SD; **** *p* value < 0.0001, *** *p* value < 0.001, ** *p* value < 0.005 compared against control(RNAi); t-test). Representative images of the expression level of the UPR^mt^ reporter are shown in the right upper panel, brightfield images in the right bottom panel.

With the aim of understanding the mechanism by which SGK-1 might regulate mitochondrial function, and possibly aging, we performed an RNAi screen. We knocked-down 836 transcription factors of the *C. elegans* genome and quantified the expression of the mitochondrial stress response reporter. We found 20 genes whose depletion reduced the expression of P*hsp-6*::GFP in *sgk-1* mutants (FC < 0.6) (Figure 2B). HIF-1 depletion did not increase P*hsp-6*::GFP (average FC 1.067) in our screen, while it did when assayed on plate (Figure 2A). One possible explanation is that liquid growing conditions are energetically more demanding, metabolism is altered and the UPR^mt^ might be differentially regulated (Lev et al., 2019). We then looked for genetic interactors by focusing on those transcription factors that when depleted reduced both the UPR^mt^ and the size of *sgk-1* mutants (Figure 2C).

Interestingly, 2 of these candidates, *lin-40* and *ceh-20*, are involved in endocytic traffic (Balklava et al., 2007). SGK-1 has been reported to be involved in endocytosis and membrane trafficking in different models (Bomberger et al., 2014; Bourgoint et al., 2018; deHart et al., 2002; Zhu et al., 2015) which supports the validity of the screen. We identified several nuclear hormone receptors (NHRs), a family of ligand-regulated transcription factors that are activated by steroid hormones, that play vital roles in the regulation of metabolism, reproduction, and development. Among them is NHR-8, a nuclear hormone receptor that regulates cholesterol levels, fatty acid desaturation and apolipoprotein production (Magner et al., 2013). NHR-8 is required for dietary restriction-mediated lifespan extension in *C. elegans* (Chamoli et al., 2014; Thondamal et al., 2014) but also normal lifespan (Magner et al., 2013). Also related with cholesterol is SBP-1, the homologue of sterol regulatory element binding protein SREBP-1. SREBPs are required to produce cholesterol (Brown and Goldstein, 1997) and regulate the expression of lipogenic genes across phyla (Horton et al., 2002; Walker et al., 2011). Interestingly, mammalian mTORC2 activates lipogenesis through SREBP-1 in the liver (Hagiwara et al., 2012). Similarly, in *C. elegans,* SBP-1 acts downstream of TOR and IIS pathways and affects adult lifespan. Depletion of *sbp-1* does not affect lifespan of wild type animals, but suppresses the increase in lifespan caused by pharmaceutical inhibition of TGF-β and IIS signaling (Admasu et al., 2018).

In summary, we identified transcription factors involved in sterol metabolism, lipogenesis and lipid membrane biology, whose depletion further reduced the size of *sgk-1* mutants (Figure 2C), indicating a genetic interaction in *C. elegans*. Combining mutations in genes that perturb the same process are expected to enhance the observed phenotypes. Loss of function of SGK-1 reduces worm size, slows down development (Jones et al., 2009; Soukas et al., 2009) and induces the UPR^mt^ (Gatsi et al., 2014). Depletion of both, NHR-8 and SBP-1, exacerbated those phenotypes in *sgk-1* mutants at the young adult stage, further reducing worm size and inducing the UPR^mt^ (Figure 2D). As reported, SBP-1 depletion reduced the size of wild type animals (Nomura et al., 2010) as did depletion of NHR-8. However, while depletion of neither NHR-8 nor SBP-1 affected development of wild type animals, it specifically slowed down the developmental rate of *sgk-1* mutants (Figure S2B), indicating a synthetic interaction. Moreover, depletion of both, NHR-8 and SBP-1, mimicked the induced UPR^mt^ observed in *sgk-1* loss-of-function mutants (Figure 2D). Apart from sterol (Roelants et al., 2018), ceramide and sphingolipids are also key structural lipids of membranes that are regulated by Ypk1/SGK-1 in yeast (Berchtold et al., 2012; Muir et al., 2014; Niles et al., 2014; Roelants et al., 2011) and regulate membrane trafficking in *C. elegans* (Zhang et al., 2011). We targeted by RNAi *sptl-1,* a serine palmitoyl-CoA acyltransferase responsible for the first committed step in *de novo* sphingolipid synthesis, and *cgt-3,* a ceramide glucosyltransferase, previously shown to interact with SGK-1 (Zhu et al., 2015). Similar to NHR-8 and SBP-1, depletion of SPTL-1 and CGT-3 induced the UPR^mt^ in otherwise wild type worms, similar to the *sgk1* mutant phenotype (Figure S2C). As reported, *cgt-3* depletion caused a very slow developmental phenotype in *sgk-1* mutants (Zhu et al., 2015). In addition, only 10-20% of animals reached adulthood, while the rest arrested development as L3 larvae. Depletion of *sptl-1* slightly slowed development and caused a partial (10 to 15%) larval arrest in otherwise wild type animals, while 100% of *sgk-1* mutants arrested development at the L3 stage. Thus, inhibition of ceramide and sphingolipids synthesis results in a synthetic interaction with *sgk-1* mutants (Figure S2D). As they are not at the same developmental stage, *hsp-6* expression levels cannot be compared, however, depletion of *sptl-1* and *cgt-3* seemed to further induce the UPR^mt^ in *sgk-1* mutants (Figure S2D). These results together, show that defective lipogenesis and/or altered cholesterol and sphingolipid metabolism cause mitochondrial proteotoxic stress. Furthermore, the results indicate that the UPR^mt^ observed in *sgk-1* mutants may be the result of altered lipid and sterol homeostasis.

### Prohibitin depletion suppresses lipogenesis and lipoprotein/yolk formation defects in *sgk-1*

SREBPs are master regulators of lipogenesis and undergo sterol-regulated export from the ER to the Golgi (Goldstein et al., 2006; Marks and Wood, 1992) where proteases release the transcriptionally active protein (Brown and Goldstein, 1997). Also, transport of ceramide from the ER to the Golgi is a crucial step in sphingolipid biosynthesis (Perry and Ridgway, 2005). Both, in *C. elegans* and mammalian models, SREBPs are regulated in response to altered membrane lipid ratios through ARF1 and GBF1 (Smulan et al., 2016), which function in vesicular trafficking, lipid homoeostasis and organelle dynamics (Kaczmarek et al., 2017). In *C. elegans* GBF-1 localizes in close proximity to the Golgi and the ER-exit sites and is required for secretion and integrity of both organelles (Ackema et al., 2013). GFB-1 and SGK-1 physically associate, GBF-1 being required for proper localization of SGK-1 near the basolateral membrane of intestinal cells (Zhu et al., 2015). Interestingly, ARF1 and GBF1 play an evolutionarily conserved role in ER-mitochondria contact site functionality and lack of GBF-1 causes a similar increase in mitochondrial size as *sgk-1* deletion in *C. elegans* ((Ackema et al., 2014) and Figure 1B and S1).

Thus, we used transmitted electron microscopy (TEM) to explore whether TORC2/SGK-1 plays a role in ER contacts and ER-mitochondria contact sites. We analyzed *sgk-1* mutants at the beginning of the reproductive period (day 1 of adulthood) and at day 5 of adulthood. We observed protruding structures from the endoplasmic reticulum, resembling exit sites, which were not observed in wild type animals, where the ER appeared compacted (Figure 3A, highlighted in Figure S3A and S3B). Lipid droplets emanate from the ER (Salo and Ikonen, 2019) and both production and secretion of lipoproteins are regulated via the ER-to-Golgi intermediary compartment (ERGIC) (Brodsky and Fisher, 2008). Protruding ER exit sites (ERES) could reflect defective lipid droplet formation and/or defective ER to Golgi or Golgi to ER secretion. TEM images showed increased number of Golgi vesicles and increased size of the Golgi system in *sgk-1* mutants (Figure 3A, highlighted in Figure S3A and S3B). The increased presence of ERES indicates impaired lipid droplet and lipoprotein biogenesis, as evidenced by the reduced content of lipid and yolk observed in *sgk-1* mutants (Figure 3A)(also visible in Figure S1A). This agrees with the role of SGK-1 in lipogenesis and vitellogenesis in *C. elegans* (Dowen et al., 2016; Wang et al., 2016; Yen et al., 2010). Interaction of lipid droplets with yolk forming particles and vacuoles also appeared altered at day 1 of adulthood and abnormal myelinated structures were visible (Figure S3B). At day 5, the number of ERES was reduced as compared to day 1 in *sgk-1* mutants, instead, small ER-emanating lipid droplets and myelinated structures/defective autolysosomes accumulate (Figure 3A)(also visible in Figure S1A). These phenotypes could reflect defective endomembrane biology in *sgk-1* mutants, in agreement with the proposed role for *sgk-1* in membrane trafficking in *C. elegans* (Zhu et al., 2015) and yeast (Clarke et al., 2017; deHart et al., 2002). In addition, the accumulation of ERES affects the proximity of ER membranes to mitochondria. In wild type animals, the ER appeared compacted and close to mitochondria, while in *sgk-1* mutants less contact between ER and mitochondria was observed in the intestine and in muscle (Figure 3A and B, respectively). This phenotype agrees with the observed localization of mTORC2 to mitochondria-associated ER membranes (MAM) and defective MAM observed in liver-specific mTORC2/rictor KO mice (Betz et al., 2013).

**Figure 3:**
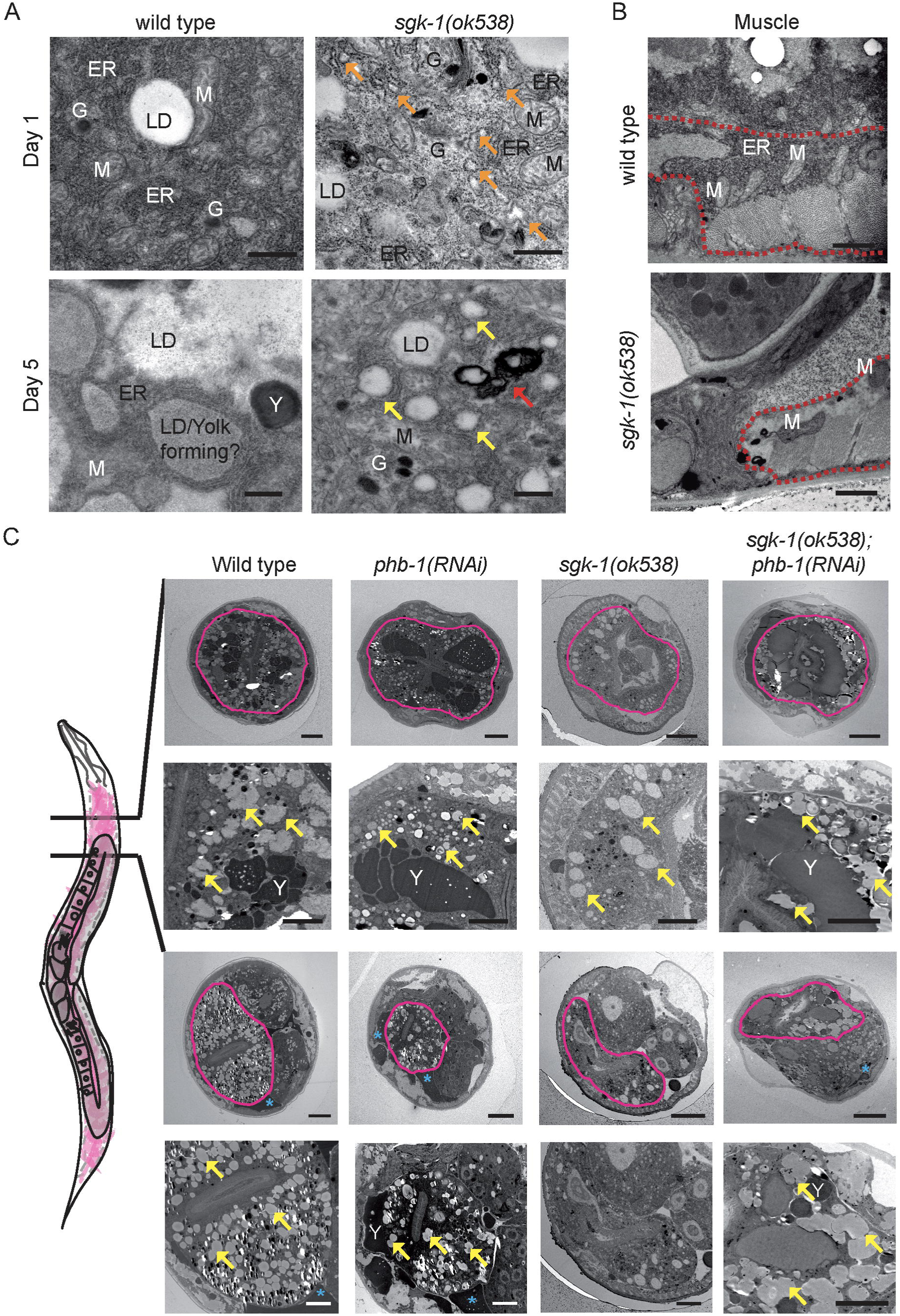
Prohibitin depletion suppresses lipogenesis and lipoprotein/yolk formation defects of *sgk-1* mutants. **A.** Transmission electron microscopy (TEM) images of the first intestinal cells of wild type and *sgk-1* mutants during aging (day 1 and day 5 of adulthood). Orange arrows point to defective ER exit sites. Yellow arrows point to forming LD. Red arrow, point to abnormal autolysosomes/myelinated forms. Bar size: 500 nm. ER: Endoplasmic Reticulum, LD: Lipid Droplet, Y: Yolk, G: Golgi. B. TEM images of muscle, delineated with a red dashed line, of wild type and *sgk-1* mutants at day 1 of adulthood. Bar size: 1 µm. M: Mitochondria C. TEM images of wild-type, *sgk-1* mutants, *phb-1* depleted animals and *sgk-1 ;phb-1 (RNAi)* treated mutants at day 5 of adulthood. Two sections, before the gonad and after the gonad turn are shown. Of each section, top images show a general view where the intestine is delineated with a pink line. Bottom images show a magnification of the intestinal area. Bar sizes: 10 µm (top panels) and 5 µm (bottom panels) for each of the cuts. Y: Yolk, blue asterisks mark pseudocoelomic lipoproteins and yellow arrows label LD.

These results support a key role of SGK-1 in membrane trafficking, most likely by affecting membrane lipid ratios and the formation of membrane contact sites, which could explain all the observed phenotypes, i.e. defective mitochondrial function and defects on lipid droplet and lipoprotein biogenesis. Since PHB depletion suppressed mitochondrial defects of *sgk-1* mutants (Figure 1), we investigated if lack of the PHB complex could suppress lipogenesis and yolk/lipoprotein accumulation. We analyzed TEM sections at the beginning of the intestine, before and after the gonad turn, in 5 days old animals (Figure 3C). *sgk-1* mutants were defective in lipid droplet formation and vitellogenesis, as previously reported (Dowen et al., 2016; Yen et al., 2010). In addition, little accumulation of lipoprotein pools at the pseudocoelom was observed compared to wild type and PHB depleted animals (Figure 3C). Strikingly, depletion of the mitochondrial PHB complex suppressed both, lipid droplet accumulation and yolk production defects of *sgk-1* mutants (Figure 3C). Furthermore, *sgk-1* mutants showed all the above described phenotypes when fed on OP50, a different bacterial food source (Figure S3C). The phenotypes observed in OP50 bacteria-fed *sgk-1* mutants included abnormal foci of myelinated forms, defective lipid droplet formation and generally reduced lipid content and yolk, as well as increased mitochondrial size.

Autophagosomes receive lipids from the ER, ERES, ERGIC, the Golgi apparatus, as well as from mitochondria and ER-mitochondria contact sites (Wei et al., 2018). Moreover, inhibition of late stages of autophagy has been shown to prevent the conversion of intestinal lipids into yolk and the accumulation of lipoprotein pools at the pseudocoelom (Ezcurra et al., 2018). Thus, our TEM analysis strongly suggests that *sgk-1* mutants suffer from a blockage of the autophagy flux.

### Prohibitin depletion suppresses the impaired autophagy flux of *sgk-1* **mutants.**

SGK-1 has been shown to modulate autophagy both via induction and blockage of autophagy (Andres-Mateos et al., 2013; Singh et al., 2013; Vlahakis et al., 2014; Vlahakis et al., 2016; Vlahakis et al., 2017; Zhou et al., 2019) and we observed an accumulation of abnormal autophagosomes in *sgk-1* mutants (Figure 3A and Figure S3B). To better understand the interaction of PHB and SGK-1, we analyzed the autophagic state in *sgk-1* mutants in the presence and in the absence of the PHB complex.

To quantify autophagy we used a strain carrying the intestinal autophagy reporter P*nhx-2*::mCherry::LGG-1 (Gosai et al., 2010) which shows a diffused cytoplasmic expression pattern under basal conditions. However, induction of autophagy results in a punctated expression pattern due to the recruitment of LGG-1 to the autophagosomes. We classified the level of autophagy based on 5 different categories from very low (animals with a completely diffused expression pattern) to very high (animals with big aggregations of LGG-1 all along the intestine) (see Materials and Methods and Figure S4A). We quantified autophagy immediately after the reproductive period (day 5), when LGG-1 puncta are more obvious. *sgk-1* mutants showed an enhanced autophagy signal compared to wild type animals (Figure 4A), as previously described in worms (Lehmann et al., 2013; Zhou et al., 2019). While depletion of the PHB complex significantly increased LGG-1 puncta in otherwise wild type worms, no additive effect was observed in *sgk-1* mutants upon *phb-1* depletion (Figure 4A). We knocked down *unc-51,* an essential gene for autophagosome formation, as a control to ensure the signal visualized in the transgenic line corresponds to autophagy and not to mCherry aggregation. *unc-51(RNAi)* reduced the autophagic signal under all tested conditions (Figure S4B).

**Figure 4:**
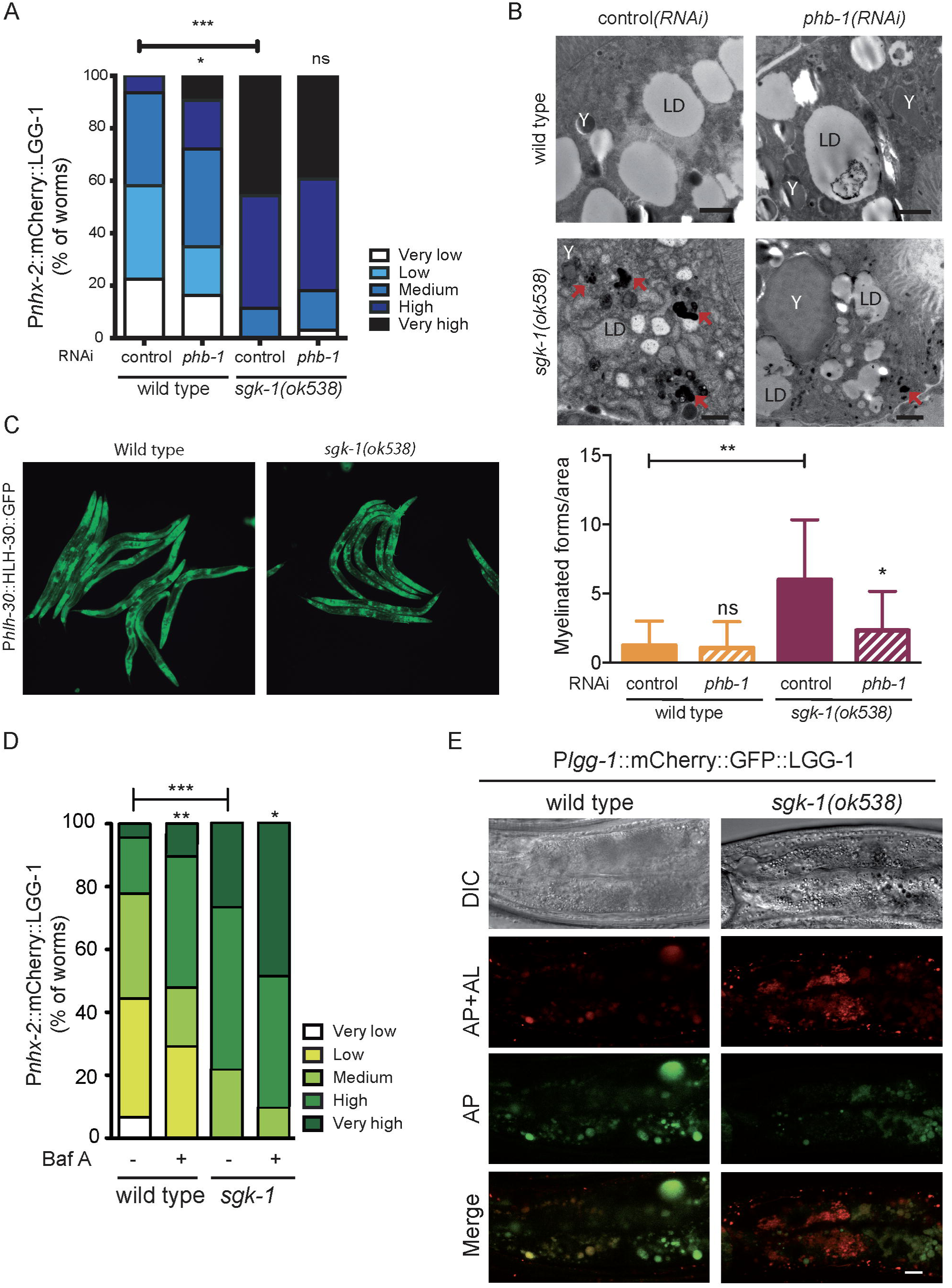
Prohibitin depletion suppresses the impaired autophagy flux of sgk-1 mutants. **A.** Quantification of the intestinal autophagy reporter Ex[Pnxh-2::mCherry::LGG-1] (see Fig. S5) in wild type animals, *phb-1* depleted worms, *sgk-1* mutants and *sgk-1 ;phb-1 (RNAi)* treated mutants at day 5 of adulthood. **B.** Electron microscopy images of the same genotypes as in A, at day 5 of adulthood. Red arrows show myelinated forms quantified in the bottom panel. LD: lipid droplet, Y: yolk particles. Bar size: 1 µm. **C.** Images of the reporter Ph/h-30::HLH-30::GFP in wild type and *sgk-1* mutants at day 1 of adulthood in OPS0. **D.** Quantification of the intestinal autophagy reporter Ex[Pnxh-2::mCherry::LGG-1] in wild type animals and *sgk-1* mutants, treated or not with bafilomycin A1 (Baf A), at day 5 of adulthood in HT115. **E.** Images of the reporter P/gg-1::mCherry::GFP::LGG-1 in wild type animals and *sgk-1* mutants at day 5 of adulthood in OPS0. Brightfield images are shown in the upper panel, red images correspond to autophagosomes (AP) and autolysosomes (AL) and green images correspond only to autophagosomes. Bottom panel show merged images. Bar size: 10 µm. (Mean ±SD; *** *p* value < 0.001,** *p* value < 0.01, * *p* value < 0.1, ns not significant; t-test. Combination of three independent replicates).

To better analyze autophagy flux, we performed electron microscopy analysis at day 5 of adulthood at the beginning of the intestine (see materials and methods). Characteristic structures of impaired autophagy flux observed in TEM images of *C. elegans* are abnormal foci with myelinated membranes (Toth et al., 2008). Myelinated forms including multivesicular bodies, multilaminar bodies and myelinated membrane whorls were clearly visible in *sgk-1(ok538)* mutants (Figure 4B). Quantification in the different backgrounds (Figure 4B, bottom panel) showed that myelinated forms rarely appeared in wild type and in *phb-1(RNAi)* treated worms, while they accumulated in *sgk-1* mutants. Interestingly, the accumulation of myelinated forms in *sgk-1* mutants was suppressed upon *phb-1(RNAi)* (Figure 4B). Altogether, the differences in LGG-1 expression pattern and the accumulation of electro-dense particles, suggest that while autophagy is induced in PHB depleted worms, both, induction of autophagy and an impairment in the autophagic flux occurs in *sgk-1* mutants, as previously proposed in yeast (Vlahakis et al., 2014; Vlahakis et al., 2016; Vlahakis et al., 2017).

We confirmed activation of autophagy in *sgk-1(ok538)* mutants by looking at the subcellular localization of the transcription factor HLH-30/TFEB, a positive regulator of autophagy, lysosome biogenesis and longevity (Lapierre et al., 2013; Settembre et al., 2011). We observed increased nuclear localization of HLH-30 in *sgk-1* mutants (Figure 4C). In order to confirm blockage of autophagy in *sgk-1* mutants we pharmacologically suppressed autophagy adding Bafilomycin A1 (BafA), an inhibitor of vacuolar type H^+^-ATPases that prevents lysosomal acidification causing a blockage of the autophagic flux (Zhang et al., 2015). Upon bafilomycin treatment, wild type worms showed an increase in LGG-1 aggregates (34%), which corresponds to the accumulation of non-degraded autophagosomes (Figure 4D). Although Bafilomycin further increased aggregates in *sgk-1* mutants (Figure 4D), this increase was considerably lower than the one observed in wild type worms (11% vs 34%), showing a partial blockage of the autophagic flux. To further confirm this blockage in *sgk-1* mutants we used the dual-fluorescent protein reporter mCherry::GFP::LGG-1, which allows monitoring of autophagosomes (APs), double-membrane vesicles that engulf cytosolic cargo, and autolysosomes (ALs), where degradation of cargo takes place after APs fuse with lysosomes (Chang et al., 2017; Kimura et al., 2007) (Figure 4E). Under the acidic environment of ALs, GFP is quenched, therefore, ALs appear as red punctae, while APs appear as yellow [green and red] punctae. In wild type animals most red puncta appeared also green, while in *sgk-1* mutants we found a large amount of only red and abnormal intestinal puncta (Figure 4E) indicating an accumulation of ALs. The same was true for the hypodermis, where *sgk-1* mutants presented more ALs than wild type worms (Figure S4C). Further demonstrating a role of SGK-1 in maintaining the autophagic flux, we observed a completely diffused expression pattern of intestinal mCherry::LGG-1 in worms overexpressing SGK-1 at day 5 of adulthood and suppression of the increased autophagosomes observed upon *phb-1* depletion (Figure S4D). Because autophagy is required for the conversion of intestinal lipids into yolk, the suppression of abnormal autophagic structures upon PHB depletion agree with the suppression of both, lipid droplet and yolk accumulation defects of *sgk-1* mutants upon PHB depletion.

### PHB depletion increases longevity in *sgk-1* mutants through modulation of autophagy and the UPR^mt^, but not mitophagy

We have shown that long-lived *sgk-1(ok538)* mutants and PHB depleted animals have induced mitochondrial quality control mechanisms such as mitophagy and the UPR^mt^ (Figure 1). Similarly, autophagy is induced in both *sgk-1* mutants and PHB depleted animals, however, the autophagy defects of *sgk-1* mutants are suppressed by PHB depletion (Figure 4). Therefore, we investigated the requirement of autophagy as well as both mitochondrial quality control mechanisms for the lifespan extension conferred by prohibitin depletion to *sgk-1* mutants. Depletion of ATFS-1 did not affect lifespan of wild type animals (Figure 5A and Table S1) (Tian et al., 2016). Surprisingly, even though prohibitin depleted worms and *sgk-1(ok538)* have induced UPR^mt^, inhibiting this process did not shorten their lifespan (Figures 5A and 5B and Table S1). However, preventing the UPR^mt^ suppressed the longevity of *sgk-1;phb-1(RNAi)* treated mutants down to wild type levels (Figure 5B and Table S1). Similarly, inhibiting autophagy by *unc-51(RNAi)* treatment did not affect lifespan of *phb-1* depleted worms nor of *sgk-1(ok538)* mutants (Figures 5C, 5D and Table S1), while it shortened that of *sgk-1;phb-1(RNAi)* treated animals (Figure 5D and table S1). Treatment with *unc-51(RNAi)* shortened the lifespan of wild type worms (Figure 5C, Table S1 and (Schiavi et al., 2013)). Autophagy is divided in four different steps (Melendez and Levine, 2009): initiation, nucleation, elongation and final fusion with lysosomes for lysis of the cargo. Likewise inhibition of initiation (*unc-51(RNAi)*), inhibition of nucleation (*bec-1(RNAi)*) and inhibition of elongation (*atg-16(RNAi)*) shortened lifespan of the *sgk-1;phb-1(RNAi)* treated mutants (Figure S5A). Therefore, we conclude that both mechanisms, autophagy and the UPR^mt^, are required for the enhanced longevity of *sgk-1(ok538)* mutants upon PHB depletion, most likely because the UPR^mt^ transcription factor is a positive regulator of autophagy.

**Figure 5:**
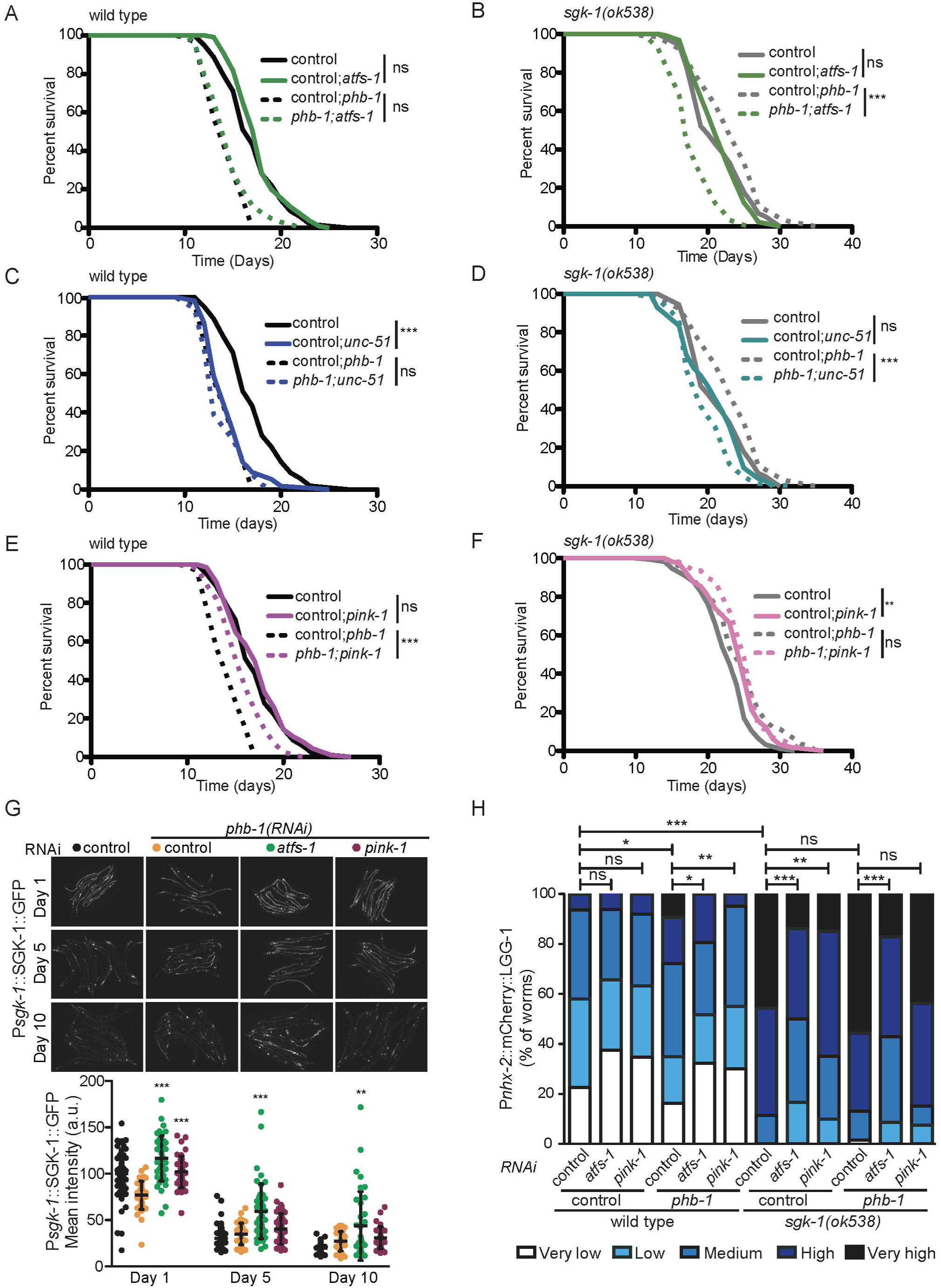
Autophagy and the UPR^mt^, but not mitophagy, are required for the enhanced longevity of *sgk-1* mutants upon PHB depletion. Lifespan of wild type worms (left) and *sgk-1* mutants (right) under normal and mitochondrial stress conditions *(phb-1(RNAi))* upon inhibition of the UPR^mt^ **(A** and **B),** inhibition of autophagy **(C** and **D)** and inhibition of mitophagy **(E** and **F).** One representative replicate is shown. Replicates and statistics are shown in Table S1. **G.** SGK-1 protein levels at day 1, day 5 and day 10 of adulthood measured using the Psgk-1::SGK-1::GFP reporter, upon PHB depletion and inhibition of either the UPR^mt^ *(atfs-1(RNAi))* or mitophagy *(pink-1(RNAi))* (Mean SD; *** p value < 0.001, ** p value < 0.01, * p value < 0.1, ns not significant;ANOVA test. Combination of at least three independent replicates is shown). **H.** Quantification of the autophagy reporter Ex[Pnxh-2::mCherry::LGG-1] in wild type animals, *phb-1(RNAi)* treated worms, *sgk-1* mutants and *sgk-1;phb-1* depleted mutants at day 5 of adulthood upon inhibition of the UPR^mt^ *(atfs-1(RNAi))* or mitophagy *(pink-1(RNAi))* (*** p value < 0.001, ** p value < 0.01, * p value < 0.1, ns not significant;t-test. Combination of at least three independent replicatesis shown).

We then assessed the requirement of mitophagy for the increased longevity of *sgk-1* mutants caused by the absence of the mitochondrial PHB complex. Inhibition of mitophagy did not affect lifespan of wild type worms (Figure 5E, Table S1 and (Palikaras et al., 2015)). Interestingly, preventing mitophagy showed a beneficial effect on lifespan of PHB depleted worms and *sgk-1* mutants (Figure 5E, 5F and table S1). However, in the case of *sgk-1;phb-1(RNAi)* treated worms inhibition of mitophagy either did not affect or increased lifespan (Figure 5F and Table S1). We confirmed the beneficial effects of preventing mitophagy in *sgk-1* and *phb-1* deficient backgrounds depleting DCT-1, a mitophagy receptor acting downstream of PINK-1 (Palikaras et al., 2015). Lack of DCT-1 increased the lifespan of PHB depleted worms and *sgk-1* mutants (Figure S5B-C and Table S1) without affecting the lifespan of wild type worms (Figure S5B and Table S1), thus recapitulating *pink-1(RNAi)* phenotypes. In the case of *sgk-1;phb-1(RNAi)* treated mutants, *dct-1* depletion increased lifespan in one out of two replicates (Figure S5C and Table S1). These results show that mitophagy can be beneficial for lifespan under certain conditions, but detrimental in certain pre-conditioned mutants such as *sgk-1* and PHB depleted worms. What determines mitophagy being beneficial or detrimental deserves further investigation.

Altogether, our results show that autophagy and the UPR^mt^ are required for the lifespan extension conferred by PHB depletion while *pink-1* mediated mitophagy is not. In agreement, in PHB depleted animals, inhibition of the UPR^mt^ resulted in a remarkable increase of intestinal SGK-1 protein levels during aging, while mitophagy inhibition did not (Figure 5G). Similar results were obtained when we pharmacologically induced mitochondrial stress with paraquat (PQ)(Figure S5D). The high SGK-1 levels observed during aging upon UPR^mt^ inhibition in combination with mitochondrial stress, could be related to the role of ATFS-1 in autophagy induction. Autophagy is one of the compensatory pathways induced to protect cells from severe mitochondrial defects (Ventura et al., 2006) and has been suggested as a cytoprotective mechanism implicated in the lifespan extension of *sgk-1* mutants upon PHB depletion (Gatsi et al., 2014). Thus, in the absence of ATFS-1, SGK-1 might become more important in keeping an efficient autophagy flux, as previously proposed (Vlahakis et al., 2016; Vlahakis et al., 2017). This hypothesis is supported by the observation that depleting genes involved in different steps of autophagy, such as initiation (*unc-51(RNAi)*) or nucleation (*bec-1(RNAi)*), increased SGK-1 protein levels (Figure S5E).

To better understand the differential requirement of both mitochondrial quality control mechanisms, UPR^mt^ and mitophagy, for the lifespan extension of *sgk-1* mutants upon PHB depletion, we examined their contribution to the observed pool of autophagosomes in *sgk-1* mutants, PHB depleted animals, and *sgk-1;phb-1(RNAi)* treated mutants (Figure 5H). When treating worms with *atfs-1(RNAi)* we observed a reduction in the autophagic signal under all tested conditions (Figure 5H), as expected since ATFS-1 regulates directly the expression of LGG-1 (Nargund et al., 2015). Treatment with *pink-1(RNAi)* reduced the accumulation of autophagosomes in *phb-1(RNAi)* and in *sgk-1* mutants, meaning that the signal observed in *phb-1(RNAi)* and in *sgk-1* mutants corresponds partially to mitophagy (Figure 5G), which agrees with the observed increase in PINK-1 protein in *phb-1(RNAi)* and *sgk-1(RNAi)* treated worms (Figure 1A). However, inhibiting mitophagy in *sgk-1;phb-1(RNAi)* did not significantly reduce the autophagosome pool (Figure 5G). Therefore, it is tempting to speculate that the mechanism by which PHB depletion extends *sgk-1* mutant lifespan is by inducing the UPR^mt^, which in turn induces general autophagy in a TORC2 independent manner.

## Discussion

Depletion of PHB, a multimeric ring-like complex sitting in the inner mitochondrial membrane induces the mitochondrial unfolded protein response (UPR^mt^) and shortens lifespan in wild type worms. In contrast, in a *sgk-1*-mutant background, PHB depletion results in a further increase in lifespan and reduced activation of the UPR^mt^ (Gatsi et al., 2014). In this work, we analyzed the interaction between the nutrient and growth factor responsive kinase SGK-1 and the mitochondrial PHB complex. Our results strongly suggest that mitochondrial, lipogenic and autophagic defects of *sgk-1* mutants are caused by defective ER function and that PHB depletion suppresses those by altering membrane lipid composition at ER-mitochondria contact sites, where TORC2 localizes, having ultimately a beneficial outcome for lifespan.

mTORC2 and SGK-1 have previously been shown to regulate mitochondrial function by maintaining mitochondria-associated ER membrane (MAM) integrity (Betz et al., 2013) and SGK-1 has been reported to localize to mitochondria upon stress (O’Keeffe et al., 2013). More recently, SGK-1 has been shown to phosphorylate VDAC1 at the outer mitochondrial membrane, inducing its degradation (Zhou et al., 2019). The data we present here supports a role for SGK-1 in MAM integrity. By electron microscopy we described bigger mitochondria in *sgk-1* mutants compared to wild type (Figure 1B). However, no obvious mitochondrial fragmentation or severe cristae defects could be observed as previously described (Zhou et al., 2019). This might be due to different growing conditions or the different alleles used. Although both predicted to be null alleles, *ok538* is a deletion mutant (Hertweck et al., 2004), while *mg455* is a nonsense mutation (Soukas et al., 2009). Nonsense mutations have been recently shown to trigger compensatory mechanisms (Wilkinson, 2019). A more detailed analysis of the different SGK-1 alleles is required to understand the different phenotypes observed. Similarly, and contrary to what has been reported (Zhou et al., 2019), we described that *sgk-1* mutants had a dramatic increase in oxygen consumption and a substantially reduced reserve capacity (Figures 1C and 1D). A low reserve capacity corresponds to the maximal usage of the mitochondrial capacity under normal conditions, which can be due to mitochondrial defects that rescind the ability of the mitochondria to function at their full potential, or to a continuous higher energy demand. Yeast Ypk1 deficient cells and *sgk-1* mutant worms have reduced mitochondrial membrane potential (Aspernig et al., 2019; Niles et al., 2014), which could indicate that a higher oxygen consumption is needed to build up a membrane potential. Our observation agrees with the increased mitochondrial respiration rate observed in mTORC2/rictor knockdown in mammalian cells (Schieke et al., 2006). Consistent with increased respiration, *sgk-1* mutants had higher ROS levels (Figure 1G and (Aspernig et al., 2019)). Interestingly, Ypk1 has been involved in both, maintenance of, and response to, ROS levels (Niles et al., 2014). Although the molecular bases of the connection between ROS levels and TORC2/Ypk1 signaling are unknown, a link with sphingolipids homeostasis has been proposed (Niles et al., 2014). The authors show that sphingolipids modulate suppression of ROS while sphingolipid depletion activates Ypk1, via ROS signaling, that in turn functions as a positive regulator of sphingolipid biosynthesis (Niles et al., 2014). Sphingolipids are essential components of biological membranes, that together with ceramides regulate intracellular trafficking, signaling and stress responses. TORC2/Ypk1 activates sphingolipid and ceramide biosynthesis (Muir et al., 2014; Roelants et al., 2011). Furthermore, sphingolipids levels modulate Ypk1 expression in a positive feedback loop to respond to membrane stress (Berchtold et al., 2012) and to regulate cell growth (Lucena et al., 2018; Roelants et al., 2018). In *C. elegans* SGK-1 has been proposed to regulate membrane trafficking through sphingolipids (Zhu et al., 2015). We show here that depletion of *sptl-1* and *cgt-3*, key genes in the synthesis of ceramides and sphingolipids, induced the UPR^mt^ as *sgk-1* deletion mutants do (Figure S2C and (Gatsi et al., 2014)) and synthetically interact with *sgk-1* (Figure S2B-D and (Zhu et al., 2015)). Interestingly, in worms, sphingolipid biosynthesis has previously been linked to the UPR^mt^, as phosphorylated sphingosine is required in mitochondria to activate the UPR^mt^ (Kim and Sieburth, 2018). Inhibition of sphingolipids biosynthesis reduces the induction of the UPR^mt^ due to a lack of ceramide, which was demonstrated by the observation that ceramide supplementation rescues *hsp-6* induction (Liu et al., 2014). Moreover, depletion of ceramide synthase genes and sphingolipids synthesis increases lifespan (Cutler et al., 2014; Menuz et al., 2009; Mosbech et al., 2013). These phenotypes resemble the effect of *sgk-1* deletion in PHB depleted animals; reduced UPR^mt^ induction and increased lifespan (Gatsi et al., 2014).

Furthermore, our transcription factor RNAi screen uncovered the implication of membrane sterol and lipid homeostasis in the UPR^mt^ induction triggered by SGK-1 deficiency (Figure 3B). Particularly interesting were NHR-8 and SBP-1/SREBP-1, considering the role of Ypk1 in sphingolipid metabolism (Berchtold et al., 2012; Muir et al., 2014; Niles et al., 2014; Roelants et al., 2011) and membrane sterol distribution at plasma membrane-endoplasmic reticulum contact sites (Roelants et al., 2018). Here we show that depletion of key transcription factors involved in lipogenesis and cholesterol metabolism, NHR-8 and SBP-1 (Horton et al., 2002; Magner et al., 2013; Walker et al., 2011), further enhances phenotypes associated with *sgk-1* mutants, including worm size reduction, developmental delay, and induction of the UPR^mt^ (Figure 3D and S2). Similarly, impaired vitellogenesis is also observed in both *nhr-8* and *sgk-1* mutants (Dowen et al., 2016; Magner et al., 2013; Wang et al., 2016). These results show that altered lipogenesis or membrane cholesterol signaling contribute to mitochondrial stress and might be common processes regulated by SGK-1, NHR-8 and SBP-1. Mammalian mTORC2 activates lipogenesis through SREBP-1 in the liver (Hagiwara et al., 2012). Therefore, it is possible that deficiencies in building up cholesterol in *sgk-1* deficient animals result in defective SREBP-1/SBP-1 activation. Moreover, cholesterol depletion prevents SGK1 signaling in *Xenopus laevis* oocytes (Krueger et al., 2009) and, in yeast, TORC2 signaling is regulated at membrane contact sites which facilitate the formation of membrane domains composed of specialized lipids (Murley et al., 2017). Our data, altogether, strongly suggest that, also in *C. elegans*, SGK-1 plays a key role in membrane lipid composition and sterol homeostasis and that organellar membrane contact sites could be modulated by TORC2/SGK-1 to coordinate cellular stress responses.

Our TEM analysis showed that *sgk-1* mutants are defective in lipid droplet and yolk/lipoprotein production, as well as pseudocelomic lipoprotein accumulation (Figure 4 and S3C). This is in concordance with the reduced lipid content and vitelogenesis observed in *sgk-1* mutants (Dowen et al., 2016; Wang et al., 2016; Yen et al., 2010) and with the proposed role of mTORC2, through SREBP1, in the regulation of lipogenesis by (Hagiwara et al., 2012). Moreover, in various systems TORC2 has been linked to functions in different membrane-contact sites (MCS) such as ER-plasma membrane (Roelants et al., 2018) and ER-mitochondria (Betz et al., 2013). Our TEM analysis supports a role for SGK-1 in ER homeostasis (Figure 4A and S3), in consonance with the induced UPR^ER^ observed in *sgk-1(ok538)* mutants (Zhu et al., 2015) and the recent uncovered role of SGK1 in the regulation of ER-dependent gene expression (Toska et al., 2019). In addition, we observed that the ER was not as close to mitochondria as in wild type animals, and in some areas in muscle cells no ER was observed at all (Figure 4B). In agreement with this, mammalian mTORC2 regulates mitochondrial physiology (Schieke et al., 2006) by localizing to mitochondria-associated ER membrane (Betz et al., 2013). Our study thus, indicates that SGK-1 plays a role in ER-mitochondria MCS that is evolutionarily conserved. The ER consists of an elaborate network of membranes that extends throughout the whole cell, making contacts with all organelles in the cell and the plasma membrane (Wu et al., 2018a). Therefore, ER defects probably transcend to all important roles of the ER in membrane biology, as we observed in *sgk-1* mutants. These include defective lipid droplet formation and lipoprotein/yolk production. Interestingly, we showed that mitochondrial PHB dysfunction suppresses defects in lipogenesis and yolk/lipoprotein production (Figure 4C). Additionally, we observed accumulation of myelinated structures/defective autolysosomes during aging in *sgk-1* mutants (Figure 4A), which suggested a blockage of the autophagic flux, most likely due to defective endomembrane interactions, as autophagosomes receive lipids from essentially all organelles and organellar-contact sites (Wei et al., 2018). Importantly, we identified a blockage of the autophagy flux in the last phase of autophagy, i.e. degradation of the cargo (Figure 4E). This agrees with yeast data where Ypk1 mutants show induced autophagy (Vlahakis et al., 2017) and reduced autophagy flux (Vlahakis et al., 2016). Thus, blockage of autophagy is conserved in SGK-1 mutants. Interestingly, depletion of PHB in *sgk-1* mutants reduced the accumulation of myelinated forms (Figure 4B). This is in line with findings in yeast, where autophagy flux is restored to wild-type levels in *ypk1* deleted *rho0* cells or upon inhibition of mitochondrial respiration under amino acid-starvation conditions (Vlahakis et al., 2016). This suggests that lack of PHB can balance membrane lipid defects of *sgk-1* mutants.

We further demonstrate that the UPR^mt^ and general autophagy, but not mitophagy, play an important role in PHB depleted *sgk-1* mutants. Autophagy and the UPR^mt^, but not mitophagy, are required for the lifespan extension conferred by PHB depletion to *sgk-1* mutants (Figure 5, S5B-C and Table S1). Similarly, inhibition of the UPR^mt^, but not mitophagy, increase SGK-1 protein levels in PHB depleted animals (Figure 5G). And finally, autophagy and the UPR^mt^/ATFS-1, but not mitophagy, contribute to the pool of autophagosomes in PHB-depleted *sgk-1* mutants (Figure 5H and S4B). We hypothesize that inhibition of the UPR^mt^ mimics inhibition of autophagy because ATFS-1 is a positive regulator of autophagy (Nargund et al., 2015). We speculate that the main function of SGK-1 is to keep organelle membrane dynamics, therefore maintaining an efficient autophagic flux. PHB depletion suppresses membrane defects of *sgk-1* mutants, making autophagy and the UPR^mt^ beneficial and essential for lifespan. In line with this, SGK-1 protein levels increase in situations of strong mitochondrial stress (*phb-1* depletion or PQ treatment) upon UPR^mt^ inhibition but not mitophagy inhibition (Figure 5G and S5D). We hypothesize that in such conditions, inhibition of autophagy by inhibiting ATFS-1, further increases the need of SGK-1 to keep an efficient autophagic flux. However, under strong mitochondrial stress, inhibition of mitophagy does not affect SGK-1 levels probably because mitophagy inhibition does not cause additional stress; instead it might be beneficial to reduce mitochondrial clearance when mitochondria are severely damaged and biogenesis is impaired. In fact, mitophagy is detrimental for both, PHB and *sgk-1* single mutants (Figures 5E, 5F and Table S1), while autophagy and the UPR^mt^ are not relevant (Figure 5B, 5D and Table S1). Recent data confirmed the beneficial effect of inhibiting mitophagy for *sgk-1* development and reproduction (Aspernig et al., 2019). Inhibition of autophagy results in increased SGK-1 protein levels (Figure S5E), further supporting a role for SGK-1 in maintenance of the autophagic flux.

We show here that the PHB complex and SGK-1 interact functionally in maintaining critical processes related to mitochondrial function, including ROS, lipid metabolism, mitochondrial size and cytosolic ROS formation, lipogenesis, yolk production and autophagy modulation. Elucidating the underlying molecular mechanisms behind the observed functional interactions between the PHB complex and SGK deserves further investigation. But many observations suggest that mitochondria associated ER membranes, MAM, might be at play. MAM has been proposed as a hub for mTORC2-Akt signaling (Betz and Hall, 2013) and TORC2/SGK1 plays a conserved role in sphingolipid and ceramide biosynthesis (Berchtold et al., 2012; Lucena et al., 2018; Muir et al., 2014; Roelants et al., 2011). The main function of MAM is to facilitate the transfer of lipids and calcium between the two organelles (Csordas et al., 1999; Rizzuto et al., 1998). MAM also mediates ER homeostasis and lipid biosynthesis by harboring chaperones and several key lipid synthesis enzymes (Stone et al., 2009; Vance, 1990), including synthesis and maturation of cholesterol, phospholipids and sphingolipids (Fujimoto et al., 2012; Hamasaki et al., 2013; Raturi and Simmen, 2013; Verfaillie et al., 2012). Besides, MAM of rat liver contains highly active sphingolipid-specific glycosyltransferases (Ardail et al., 2003), and ceramide metabolism has been proposed to occur at mitochondria as well as at MAM (Bionda et al., 2004). The PHB complex affects membrane lipid composition and genetically interacts with proteins involved in ER-mitochondria communication. PHB has been proposed to play a key role as membrane organizer since it affects the distribution of cardiolipin and phosphatidylethanolamine (Birner et al., 2003; Klecker et al., 2015; Osman et al., 2009). In addition, lack of PHB in MEFs affects lipid metabolism and cholesterol biosynthesis (Richter-Dennerlein et al., 2014). Moreover, PHB proteins have been genetically linked to the mitochondrial inner membrane organizing system (MINOS) and to ER-mitochondria encountered structures (ERMES) complexes in synthetic lethal screens in yeast, forming part of a large ER-mitochondria organizing network, ERMIONE (Birner et al., 2003; Kornmann et al., 2009; van der Laan et al., 2012). On the other hand, PHB has been related with sphingosine metabolism as it interacts with sphigosine-1-phosphate in mice mitochondria (Strub et al., 2011).

In the future, it will be important to decipher the role of SGK-1 at MAM and MCS at the molecular level and the mechanism by which PHB depletion suppresses *sgk-1* defects. We hypothesize an imbalance between PHB and SGK-1 activity caused by depletion of either one has serious consequences for mitochondrial maintenance and cellular homeostasis. Removing both restores balance. By studying the interaction of PHB with TORC2/SGK-1 we unravel a conserved key role for both components in lipid homeostasis. Our data demonstrates that PHB deficiency could alter membrane lipid composition in a favorable manner for *sgk-1* mutants. Given the implication of TORC2, PHB and autophagy in aging and cancer, our findings may contribute to ameliorate the pathogenesis of aging-related disorders.

## Materials and Methods

### *C. elegans* strains and maintenance

The *C. elegans* strains used in this study were: N2 (wild type); MRS486: *sgk-1(ok538)X* (3x outcrossed from BR4774, in total 11x outcrossed to N2 from the CGC strain VC345; MRS88: *daf-2(e1370)III;* MRS68: *daf-2(e1370)III*;*sgk-1(ok538)X;* MRS386: *byEx*[*Psgk-1::*SGK-1::GFP *+ rol-6(su1006)* (1x outcrossed from BR2773 (Hertweck et al., 2004)); MRS38: *daf-2(e1370)III*;*byEx[*P*sgk-1::*SGK-1::GFP *+ rol-6(su1006)];* VK1093: *vkEx1093[*P*nhx-2::*mCherry::LGG1]; MRS92: *sgk-1(ok538)X;vkEx1093[*P*nhx-2::*mCherry::LGG1]; MRS112: *byEx[*P*sgk-1::*SGK-1::GFP *+ rol-6(su1006)]; vkEx1093[*P*nhx-2::*mCherry::LGG1]; SJ4100: *zcIs13[*P*hsp-6::*GFP*]V;* MRS19: *sgk-1(ok538)X*;*zcIs13[*P*hsp-6::*GFP*]V;* MRS63: *rict-1(ft7)II*;*zcIs13[*P*hsp-6::*GFP*]V*; BR4006: *pink-1(tm1779)II;byEx655[*P*pink-1::*PINK-1::GFP *+* P*myo-2::*mCherry + herring sperm DNA]; MAH215: *sqIs11*[P*lgg-1*::mCherry::GFP::LGG-1 + *rol-6(su1006)*]; MRS555: *sgk-1(ok538)X;sqIs11*[P*lgg-1*::mCherry::GFP::LGG-1 + *rol-6(su1006)*]; MAH240: *sqIs17*[P*hlh-30*::HLH-30::GFP + *rol-6(su1006)*]; MRS558: *sgk-1(ok538)X;sqIs17*[P*hlh-30*::HLH-30::GFP + *rol-6(su1006)*]; MRS505: *sod-1(tm776)II* (2x outcrossed to N2); MRS516: *sod-1(tm776)II;sgk-1(ok538)X*; MRS504: *sod-2(gk257)I* (2x outcrossed to N2); MRS510: *sod-2(gk257)I;sgk-1(ok538)X*.

Unless otherwise stated, we cultured the worms according to standard methods (Brenner, 1974). We maintained nematodes at 20°C on nematode growth media (NGM) agar plates seeded with live *Escherichia coli* OP50 (obtained from the CGC).

### RNAi assays on plates

For RNAi experiments worms were placed on NGM plates, supplemented with 25 µg/ml carbenicillin (Sigma-Aldrich) and 10 µg/ml nystatin (Sigma-Aldrich), seeded with HT115 (DE3) *Escherichia coli* bacteria (deficient for RNase-E) transformed with empty vector (control) or the required target gene RNAi construct. Each bacterial strain was inoculated, from an overnight pre-inoculum, in LB (ampicillin (100 µg/ml) (Sigma-Aldrich) and tetracycline (15 µg/ml) (Sigma-Aldrich). The overnight culture was diluted (1:10) and grown in a shaking incubator at 37°C for 3 hours, until it reached an OD_600_ of around 1.5. Then, in order to induce the dsRNA expression, we added IPTG (final concentration: 1 mM) (Sigma-Aldrich) to the bacterial culture and we harvested the bacterial culture after 2 hours of incubation at 37°C by centrifugation (8 min at 6000×*g,* 4°C). Following, we washed the pellets with S Basal and harvested them again. Finally, we re-suspended bacterial pellets to a final concentration of 30 g/l in complete S Medium. Bacterial stocks were kept at 4°C up to 4 days before being used. For all the RNAi treatments, bacterial stocks were mixed in a proportion of 1:1 before seeding the plates.

A semi-synchronous embryo population was grown on plates seeded with the appropriate RNAi bacterial clone at 20°C until the desired stage (young adult, day 1, day 5 or day 10 of adulthood). During the egg-laying period, we transferred worms every day and every 2 days thereafter.

To pharmacologically induce a strong mitochondrial stress, we transferred L3 to fresh RNAi plates containing 0.25 mM Paraquat (Sigma-Aldrich).

### Imaging

On the day of imaging, worms were anesthetized with 10 mM Levamisole (Sigma-Aldrich) or with a mixture of 10 mM Levamisole and 5 mM NaN_3_ (Sigma-Aldrich) for confocal imaging and mounted on 2% agarose pads. P*pink-1::*PINK-1::GFP, P*nhx-2::*mCherry::LGG1 and P*sgk-1::*SGK-1::GFP reporter were observed using the AxioCam MRm camera on a Zeiss ApoTome Microscope; P*lgg-1*::mCherry::GFP::LGG-1 reporter was imaged with a confocal Nikon A1R equipped with a Plan Apo VC 60x/1.4 objective and P*hsp-6::*GFP animals were visualized with a Leica M205 Stereoscope equipped with a Plan Apo 5.0x/0.50 LWD objective and a ORCA-Flash4.0 LT Hamamatsu digital camera.

Image analysis was performed using the ImageJ software. Emission intensity was measured on greyscale images with a pixel depth of 16 bits. At least two independent assays were carried out and the combined data was analyzed using the GraphPad Prism software (version 5.0a).

### Electron microscopy

Transmission electron microscopy (TEM) of *C. elegans* was carried out as described (Hall et al., 2012) with small modifications. Adult worms, at day 1 or 5, were immersed in 0.8% glutaraldehyde + 0.8% osmium tetroxide in 0.1 M sodium cacodylate buffer, pH7.4. Carefully, under a dissecting scope, we cut the animals with a scalp at the posterior end of the intestine, removing the tail, and kept the samples for 1 hour on ice under dark conditions to allow complete penetration of the fixing solution. We washed the worms three times with 0.1 M sodium cacodylate buffer and fixed them over night at 4°C in 2% osmium tetroxide in 0.1 M sodium cacodylate buffer; the entire procedure was performed on ice. On the next day, we washed the worms three times with 0.1 M sodium cacodylate buffer on ice and finally embedded the worms in small cubes of 1% agarose. We then dehydrated the samples by incubating them in 50%, 70% and 90% ethanol for 10 minutes each, followed by 3 washes of 10 minutes each in 100% ethanol, at room temperature. Next, we incubated the samples in a 50:50 ethanol/propylene oxide solution for 15 minutes followed by 2 incubations of 15 minutes each in 100% propylene oxide. We then infiltrated the samples on a rotator or with an embedding machine for 2 hours in a 3:1 ratio of propylene oxide to resin and next for 2 hours in a 1:3 ratio of propylene oxide to 3 resin. Following, we incubated the samples overnight in 100% resin and the next day changed to fresh 100% resin for 4 hours. We finally arranged the samples in a flat embedding mold and cured them at 60°C for 2 days.

We cut the worms at the level of the first intestinal cells, immediately before and after the gonad turn, specifically at 175 µm, 185 µm, 250 µm and 350 µm from the mouth.

For myelinated forms quantification, we analyzed 12 areas of 56 μm^2^ per condition and counted together multivesicular bodies (MVB), multillaminar bodies (MLB) and myelinated forms. Data was analyzed using the GraphPad Prism software (version 5.0a).

### Oxygen consumption

Worm oxygen consumption was measured using the Seahorse XFp Analyzer (Agilent) as described (Luz et al., 2015). Briefly, worms at the young adults stage (30 per well) or day 6 adult worms (20 per well), were transferred into M9-filled Seahorse plates. OCR was measured 8 times in basal conditions as well as after each injection. Working solutions were diluted in M9 at the final concentrations: FCCP (Sigma-Aldrich) 250 μM, oligomycin (Sigma-Aldrich) 250 μM and NaN_3_ (Sigma-Aldrich) 400 mM. Data were normalized by the protein content quantified by the bicinchoninic acid assay (Thermofsiher). The OCR values were arranged in order to see the response to the drug. Basal OCR corresponded to the average of the 8 measurements under basal conditions, while the rest of the values derived from the average of the last 4 measurements. At least three independent assays were carried out and the combined data was analyzed by *t*-test using GraphPad Prism software (version 5.0a).

### Quantification of Reactive oxygen species

Cytoplasmic ROS was quantified using 2,7-dichlorofluorescin-diacetate (H2-DCFDA) dye (Sigma-Aldrich) modified from (Artal-Sanz and Tavernarakis, 2009b). Briefly, 150 nematodes at the young adult stage were manually selected and recovered in M9 buffer to a final concentration of 1 worm/μl. 50 μl of the worm suspension was transferred into a 96-well flat bottom plate in triplicate (Thermo Fisher Scientific) and 50 μl of 100uM H2-DCF-DA solution was added to the wells (H2-DCF-DA to a final concentration of 50 μM). Immediately, basal fluorescence was measured (T_o_) (Ex/Em: 485/520 nm) using a microplate reader (PolarStar Omega. BMG, LabTech). After 1 hour of incubation at 20 °C shaking, fluorescence was measure one again (T_1_). The final fluorescence was calculated by T_1_-T_0_. Data were normalized by the protein content quantified by the bicinchoninic acid assay (BCA) (Thermo Fisher Scientific) and manually counting the number of worms per well. At least three independent assays were carried out and the combined data was analyzed by *t*-test using GraphPad Prism software (version 5.0a).

### Protein Content Quantification

Total protein content was determined using the bicinchoninic acid (BCA) method. Briefly, pellets from 50 worms were dried in a Speed Vac Concentrator (SPD12 1P SpeedVac, Thermo Scientific), 20 µl of 1 M NaOH were added to the dry pellet. Fat was degraded by heating at 70°C for 25 min and 180 µl of distilled water was added.

After vortexing, the tubes were centrifuged at 14000 rpm for 5 min and 25 µl of the supernatant were transferred into a 96 well plate. Next, 200 µl of the BCA reagent, prepared according to manufacturer’s instructions (Pierce BCA Protein Assay Kit, Thermo Scientific) was added to the sample. After incubation at 37°C for 30 min, the plate was cooled to room temperature and absorbance was measured using the POLARstar Omega luminometer (BGM Labtech) at 560 nm.

### Autophagy assays

In order to monitor autophagy in *C. elegans* we used the reporter strain Ex[*Pnhx-2*::mCherry::LGG-1] (Gosai et al., 2010). Animals were classified in 5 different expression categories (Figure S4A): <u>Very low:</u> animals with a completely diffused expression pattern; <u>Low:</u> animals with a generally diffuse expression pattern but with the presence of small puncta; <u>Medium:</u> animals with puncta along the intestine; <u>High:</u> animals showing aggregation of LGG-1 puncta mostly in the posterior part of the intestine; <u>Very high:</u> animals with large LGG-1 aggregates along the whole intestine. To avoid bias in the classification process, different researchers performed blind scoring on the same animal images and disputes were resolved after discussion.

To determine whether increased levels of LGG-1 puncta corresponded to induced autophagy or a blockage in the flux, we treated day 5 adult worms with Bafilomycine A_1_(BafA; Sigma-Aldrich), which blocks lysosome acidification. Animals were washed from plates with M9 buffer and harvested in 15 ml tubes. In order to remove excess of bacteria, we washed animals at least three times with M9 by letting the worms to settle at the bottom of the tubes and then removed the supernatant. After washes, worms were incubated for 6 hours in M9 with BafA (1.6 mM) or DMSO (1.5%) at 20°C with agitation.

At least 3 independent assays were carried out and the combined data was analyzed by *t-test* using GraphPad Prism software (version 5.0a).

### Lifespan Analysis

All lifespans were performed at 20°C. Semi-synchronized eggs were obtained by hypochlorite treatment of adult hermaphrodites and placed on NGM plates seeded with HT115 *E. coli* bacteria. During the course of the lifespan assays, we transferred adult worms to fresh plates every day during their reproductive period and afterwards on alternate days. Worms were scored as dead when they no longer responded to touch, while exploded animals, those exhibiting bagging (embryo hatching inside the worm), or dried out at the edges of the plates were censored.

For N-Acetyl-L-Cysteine (NAC) (Sigma-Aldrich) treatments we added NAC to the RNAi medium to a final concentration of 8 mM.

We used the GraphPad Prism software (version 5.0a) to plot survival curves and to determine significant differences in lifespans (log-rank (Mantel-Cox) test). See Table S1 for lifespan statistics.

### RNAi screen

We performed an image-based RNAi screen as previously described(Hernando-Rodriguez et al., 2018). Briefly, we dispensed synchronized *sgk-1(ok538);*P*hsp-6*::GFP L1 larvae in 96 well plates using a microplate dispenser (EL406 washer dispenser, BioTek) and added the corresponding dsRNA encoding bacterial culture. Next, we incubated the worms at 20°C with constant shaking (120 rpm - New Brunswick™ Innova^®^ 44/44R) until they reached the young adult stage. We acquired pictures of each well using the IN Cell Analyzer 2000 (GE Healthcare) and performed the image analysis with a user-defined protocol which was developed on the Developer Toolbox software (GE Healthcare). We tested 836 RNAi clones in duplicate and candidates were defined based on the adjusted *p* value and the fold change (FC) (*P* value < 0.001 and FC < 0.6).

## Acknowledgements

Some strains were provided by the Caenorhabditis Genetics Center (CGC), which is funded by NIH Office of Research Infrastructure Programs (P40 OD010440). Special thanks to Ralf Baumeister (Albert-Ludwig University, Germany) for the strain BR2773:*byEx*[*Psgk-1::*SGK-1::GFP] (Hertweck et al., 2004) and to Nuria Flames for sharing her transcription factor RNAi library. Thanks to Mario Soriano Navarro, from the CIPF, for technical help with TEM. M.A-S was supported by the Ramón y Cajal program of the Spanish Ministerio de Economía y Competitividad (MINECO), RYC-2010-06167. B.H.R. was supported by the Spanish MINECO FPI program, BES-2013-064047. Our work is supported by the Spanish Ministerio de Economía y Competitividad (BFU2012-35509), and the European Research Council (ERC-2011-StG-281691).

## Author contributions

B.H.R., M.M.P.J. and M.A.-S. designed the experiments; B.H.R., M.M.P.J., M.J.R.-P. A.P., M.D.M.-B. P.dl.C.R., R.G. and M.A.-S. carried out experiments; B.H.R., M.M.P.J., M.J.R.-P. A.P., M.D.M.-B. P.dl.C.R., R.G. and M.A.-S analyzed the data and interpreted results and B.H.R. and M.A.-S wrote the manuscript. All authors read, commented and approved the final manuscript.

